# Salt-inducible kinase inhibition promotes weight loss and improves the diastolic function of obesity-related heart failure with preserved ejection fraction in mice

**DOI:** 10.1101/2025.03.02.640935

**Authors:** Fubiao Shi, Vineet Agrawal, Uugantsetseg Munkhjargal, Elizabeth Kobeck, Hari U. Patel, Wei Zhang, Sheila Collins

**Affiliations:** Division of Cardiovascular Medicine, Department of Medicine, Vanderbilt University Medical Center, Nashville, TN 37232, USA; Tennessee Valley Healthcare System Nashville Veteran Affairs Hospital, Nashville, TN 37212, USA; Department of Molecular Physiology and Biophysics, Vanderbilt University School of Medicine, Nashville, TN 37232, USA

## Abstract

**Background:** Obesity negatively impacts cardiac function and is closely associated with heart failure with preserved ejection fraction (HFpEF). Salt-inducible kinases (SIKs) are critical regulators of energy metabolism and cardiovascular function. Small molecule SIK inhibitors have been developed but their effect in treating obesity-related HFpEF remains unexplored. We recently discovered that pharmacological SIK inhibition promotes the adipose tissue thermogenesis and mitochondrial biogenesis gene program. We reason that targeting SIKs to treat obesity-related complications would be beneficial for HFpEF.

**Methods:** We employed a preclinical HFpEF mouse model induced by two-hit stress of high-fat diet (HFD) and nitric oxide synthase (NOS) inhibition using L-NAME in drinking water. Eight-week-old C57BL/6J mice received a regular low-fat chow diet or HFD/L-NAME for 5 weeks and were treated with vehicle or a pan-SIK inhibitor YKL-05-099 (YKL) via daily intraperitoneal injection in the last 4 weeks. Body weight and parameters of adiposity, energy balance and glucose tolerance were assessed. Cardiovascular function was characterized by echocardiography and *in vivo* pressure-volume loop hemodynamic analysis. Myocardial transcriptomic data were analyzed to determine if changes in SIK gene expression are associated with human HFpEF.

**Results:** YKL treatment limits body weight gain mainly by reducing the fat mass in obese HFpEF mice. YKL-treated mice show better glucose tolerance, enhanced adipose tissue browning and decreased lipid deposition. YKL-treatment mice demonstrate preserved left ventricular (LV) ejection fraction, reduced LV filling pressure and improved diastolic function. Myocardial expression of *SIK1, SIK2*, and *SIK3* mRNA is down-regulated in patients with HFpEF. However, higher SIK mRNA expression is associated with a subgroup of HFpEF patients that has a greater risk for HF hospitalization and/or death.

**Conclusions:** Taken together, our study reveals a pathological role for SIKs in obesity-related HFpEF and suggests that pharmacological SIK inhibition would be a disease-modifying strategy for obese HFpEF, for which evidence-based therapy has been limited.

## Introduction

Heart failure with preserved ejection fraction (HFpEF) is a prevailing and heterogenous condition. It is defined as HF with left ventricular ejection fraction (LVEF) ≥ 50% and elevated LV filling pressures at rest or during exercise^1^. It accounts for approximately half of hospital admissions of individual with HF^2^ and carries a 75% five-year mortality rate^3^, but evidence-based therapy has been very limited^4^. The complex clinical features of HFpEF include the presence of multiple comorbidities, including hypertension, obesity, diabetes, and metabolic dysfunction^5^. Coincidence of metabolic and hypertensive stress, induced by high-fat diet (HFD) and endothelial nitroxide synthases (eNOS) inhibition using Nω-nitrol-arginine methyl ester (L-NAME), leads to systemic and cardiovascular features of human HFpEF in mice^6^, suggesting obesity-induced metabolic dysfunction is a critical pathological driver in HFpEF.

As an energetically demanding organ, the healthy adult heart has a high metabolic flexibility and predominantly (∼95%) relies on mitochondrial oxidative metabolism for ATP production^7^. In contrast, the failing heart is subjected to a shift of substrate reliance from oxidative to glycolytic metabolism, primarily due to compromised mitochondrial function and decreased oxidative metabolism^7^. For instance, metabolomic and transcriptomic studies have identified circulating and myocardial signatures indicative of mitochondrial dysfunction, impaired fatty acid utilization and myocardial fuel inflexibility in human HFpEF^8-10^. Metabolic dysfunction results in abnormal myocardial active relaxation and an increase in passive stiffness due to metabolic reprograming and structural remodeling and ultimately leads to diastolic dysfunction^11,12^.

Salt-inducible kinases (SIKs), including SIK1, SIK2 and SIK3, are serine/threonine kinases of the AMP-activated protein kinase (AMPK)-related kinase family^13^. Protein kinase A (PKA)-mediated SIK inhibition is a major link of G-protein coupled receptor activation and the downstream target gene transcription program^14,15^. Previous studies including ours have shown that SIKs are downstream mediators of the β-adrenergic receptor signaling pathway and function as a suppressor of the adipocyte thermogenic gene program^16-20^. Our study further showed that pharmacological and genetical inactivation of SIKs in brown adipocytes promotes the thermogenic and mitochondrial gene program^20^. In mice, SIK inhibition promotes the expression of key thermogenic protein uncoupling protein 1 (*Ucp1*) in the white adipose tissue^20^. Here we show that treatment with a pan-SIK inhibitor promotes weight loss and improves diastolic function in the L-NAME/HFD-induced obese HFpEF mouse model. Our data suggest that pharmacological SIK inhibition would be a potential disease-modifying strategy for obese HFpEF, for which evidence-based therapy has been limited^4^.

## Results

### YKL treatment reduce bodyweight and limit adiposity in obese HFpEF mice

Our previous study showed that treatment with a pan-SIK inhibitor YKL-05-099^21^ (YKL) increases adipose tissue browning^20^, suggesting SIK inhibitor could be a weight loss reagent in mice. As it has been well-appreciated that obesity-induced metabolic dysfunction is a critical driver for HFpEF, we hypothesize that pharmacological SIK inhibition will prevent obesity-related complications and thereby improve the cardiometabolic outcomes of obese HFpEF. To test this hypothesis, we employed a preclinical two-hit obese HFpEF mouse model induced by HFD and eNOS inhibition using L-NMAE^6^. To induce the HFpEF phenotype, eight-week-old male C57BL/6J mice were given L-NAME in drinking water and fed with HFD for 5 weeks (**Figure 1A**). To probe the therapeutic effects of SIK inhibitor, obese HFpEF mice were treated with vehicle or YKL (10 mg/kg bodyweight) by daily intraperitoneal injection for 4 weeks from week 2 to 5 of L-NAME/HFD treatment, and their metabolic and cardiac function were characterized (**Figure 1A**). C57BL/6J mice were fed with low-fat chow diet (LFD) for 5 weeks and injected with vehicle in the last 4 week to serve as control. After 4-week YKL treatment, body weight of the treatment group is significantly lower, mainly due to reduced fat mass without loss of lean mass (**Figure 1B**). The weight of inguinal and gonadal white adipose tissues (iWAT and gWAT) was significantly reduced, and the weight of brown adipose tissue (BAT), liver, quadriceps muscle and kidney does not change (**Figure 1C**). Compared with baseline, the gained body weight of the YKL treatment group is significantly reduced (**Figure 1D**). Daily food intake of the YKL treatment group does not change (**Figure 1E**), but their daily fecal calorie content trends to be decreased (**Figure 1F**), suggesting a potential negative energy balance in obese HFpEF mice after YKL treatment. In addition, glucose tolerance was also moderately improved in the YKL-treatment mice, as indicated by a trend of faster glucose clearance post a glucose bolus injection (**Figure 1G**). Taken together, our data shows that YKL treatment reduces adiposity and improves glucose homeostasis in obese HFpEF mice.

**Figure 1.**
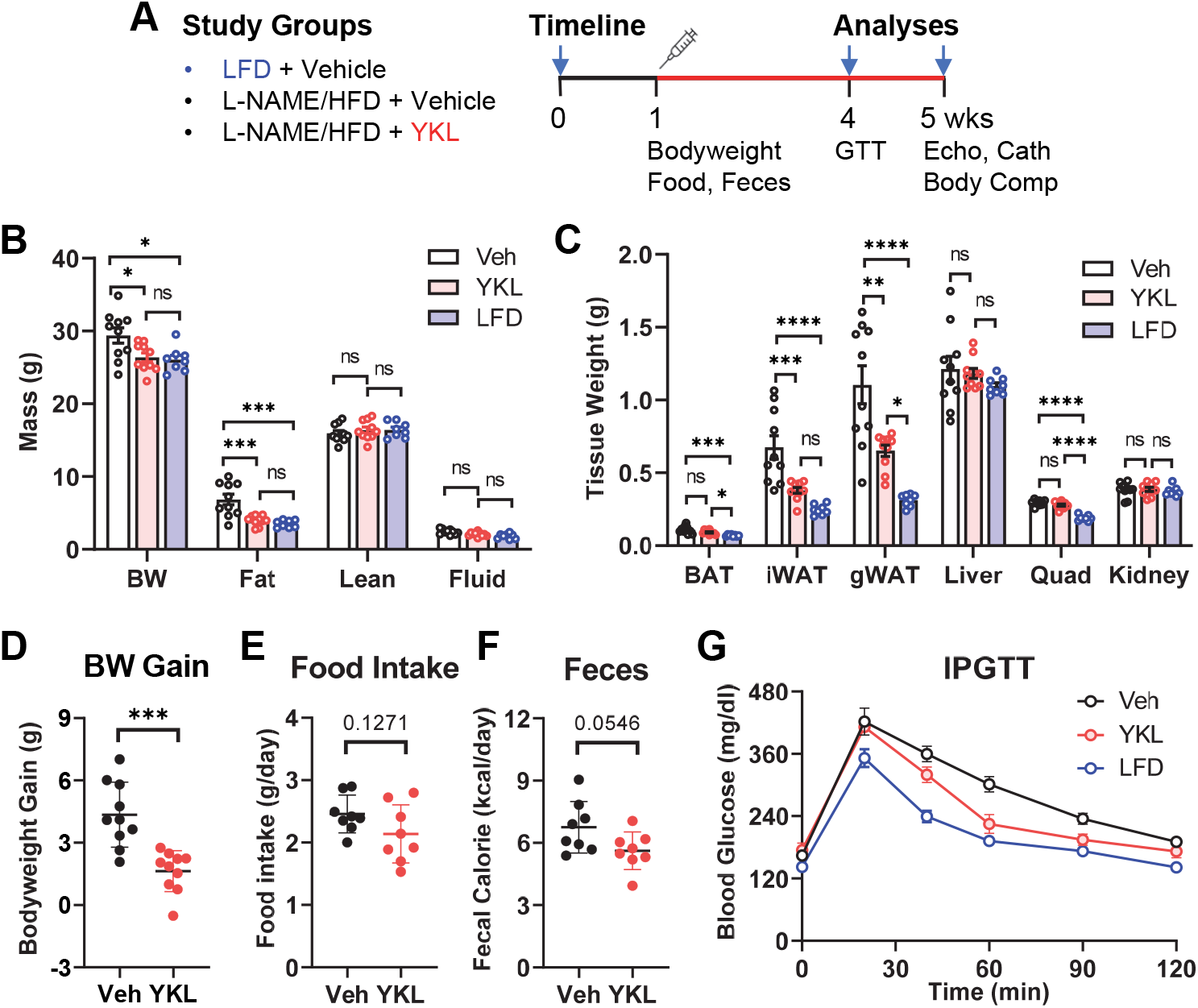
SIK inhibition reduces body weight and improves glucose tolerance in obese HFpEF mice. (A) Study groups and timeline. Eight-week-old C57BL/6J were fed with LFD or HFD/L-NAME for 5 weeks. Mice were administrated vehicle or YKL (10 mg/kg) via daily intraperitoneal injection for 4 weeks. Analyses of the metabolic and cardiac function were performed in week 4 and 5. GTT: glucose tolerance test. Echo: echocardiography. Cath: cardiac catheterization. Body Comp: body composition. (B-C) Body weight (BW), body composition (fat, lean and fluid mass), tissue weight at the end of week 5. BAT: brown adipose tissue. iWAT: inguinal white adipose tissue. gWAT: gonadal WAT. Quad: quadriceps muscle. (D-F) Body weight gain, daily food intake, and daily fecal calorie contents. (G) Intraperitoneal glucose tolerance test (IPGTT) at the end of week 4. P-values of one-way ANOVA (B-C) and t-tests (D-F). *p<0.05, **p<0.01, ***p<0.001. ns: not statistically significance.

### YKL treatment promotes adipose tissue browning and reduces hepatic lipid deposition

To determine if YKL treatment has a direct impact on adipose tissues, we next examined adipose tissues by histology. Hematoxylin and eosin (H&E) staining shows that adipocytes with multilocular lipid droplets appear in the iWAT of YKL treated obese HFpEF mice, while the lipid droplets in their BAT are much less (**Figure 2A**). Immunohistochemistry staining shows that the key thermogenic protein uncoupling protein 1 (UCP1) is significantly induced in the iWAT of YKL-treated mice (**Figure 2B**), suggesting enhanced adipose tissue browning in response to YKL treatment. In addition, although total liver weight does not change (**Figure 1C**), hepatic lipid droplets was much less in the YKL-treated mice (**Figure 2C**). Quantification of H&E-stained tissue sections showed that the lipid contents are significantly reduced both in BAT and liver of YKL-treated mice (**Figure 2D**). These results suggests that YKL treatment promotes adipose tissue thermogenesis and reduces lipid deposition, further supporting an adiposity-reducing effect in obese HFpEF.

**Figure 2.**
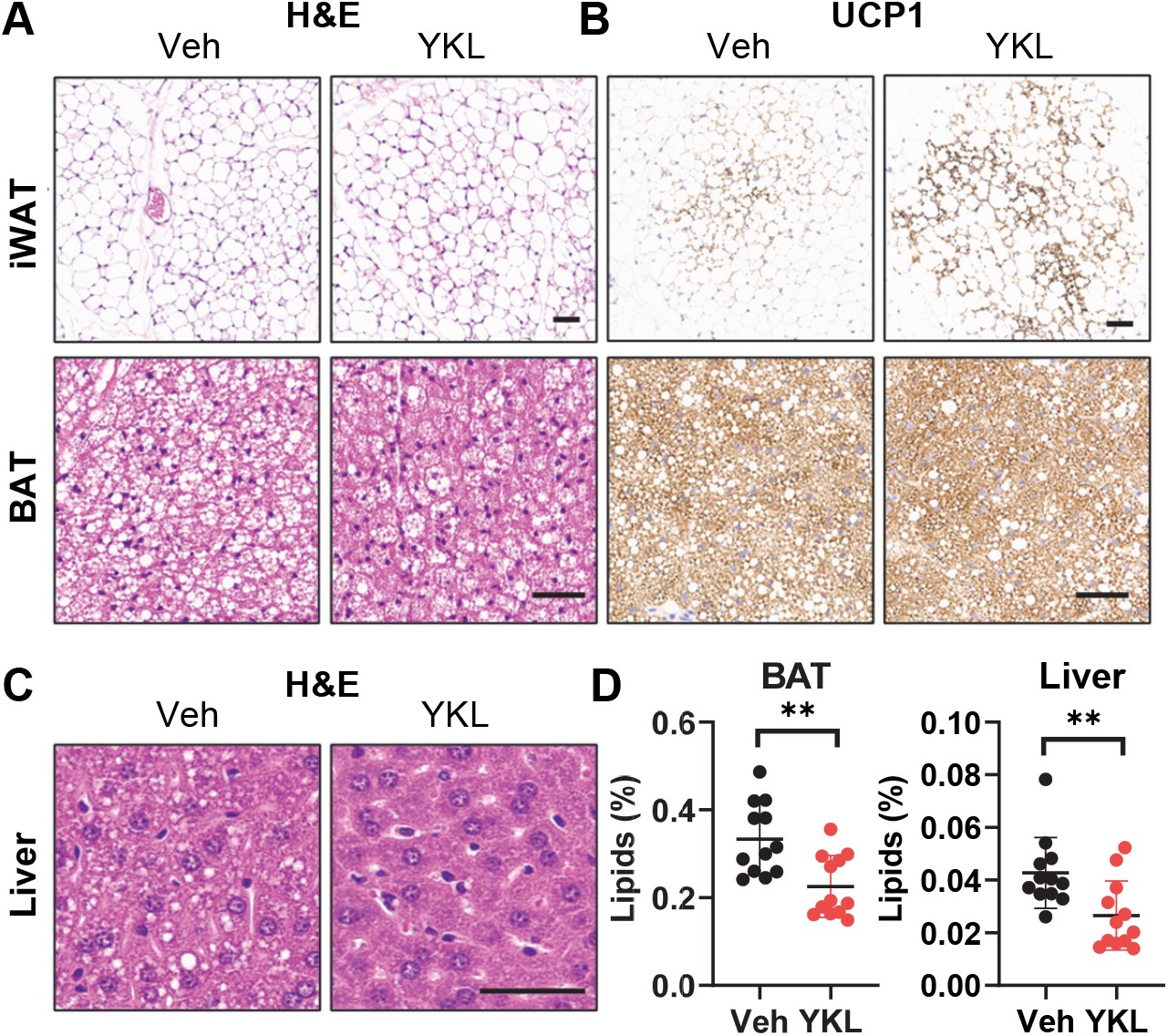
SIK inhibition promotes adipose tissue browning and reduces lipid deposition in obese HFpEF mice. (A-B) Hematoxylin and eosin (H&E) and UCP1 immunohistochemical staining of iWAT and BAT sections from mice after 4-weeks of vehicle (Veh) and YKL treatment. (C) H&E staining of liver sections from mice after 4-weeks of treatment. (D) Qualification of lipid contents on H&E-stained BAT and liver sections. Scale bar: 50μM in A-C. P-values of Mann-Whitney test (D). **p<0.01.

### YKL treatment improves diastolic function of obese HFpEF mice

Our data shows that YKL treatment does not alter the left ventricular (LV) and right ventricular (RV) weights of obese HFpEF mice (**Figure 3A**). To determine if YKL treatment affects the cardiac function of obese HFpEF mice, we next performed echocardiography and cardiac catheterization for *in vivo* pressure-volume loop hemodynamic measurements, metrics that are used in humans to diagnose and monitor diastolic function. Echocardiography shows that the LVEF is preserved, and the cardiac output and heart rates remains comparable to vehicle-treated obese HFpEF mice (**Figure 3B**). However, YKL treatment leads to significant morphological alterations in the heart, including reduced end-diastolic left ventricular posterior wall thickness (LVPWd) and interventricular septum thickness (IVSd) (**Figure 3C**). In addition, the left atrial (LA) diameter and LA stroke volumes of end-systolic and end-diastolic status are also significantly reduced (**Figure 3D**). Furthermore, non-invasive Doppler imaging shows that LV filling pressures are decreased in the YKL-treated mice, as indicated by reduced ratio of early mitral wave velocity and late mitral wave peak velocity (E/A) and reduced ratio of early mitral wave velocity and early diastolic mitral annular velocity (E/e’) (**Figure 3E**). Invasive *in vivo* hemodynamic measurement further confirmed a robust decrease in LV systolic and diastolic pressures, and diastolic time constant (Tau) (**Figure 3F**). Furthermore, the RV systolic and diastolic pressures, and diastolic time constant (Tau) are also significantly decreased in YKL-treated mice (**Figure 3G**). Of note, some of morphological and functional alterations in YKL treatment mice are almost normalized to an extent that is comparable to LFD control mice (LVPWd, IVSd, LA diameter, and E/A), suggesting a direct disease modifying effect in obese HFpEF hearts. Taken together, echocardiography and hemodynamic measurements demonstrates a direct treatment effect of YKL in the heart that indicates improved diastolic function in obese HFpEF mice.

**Figure 3.**
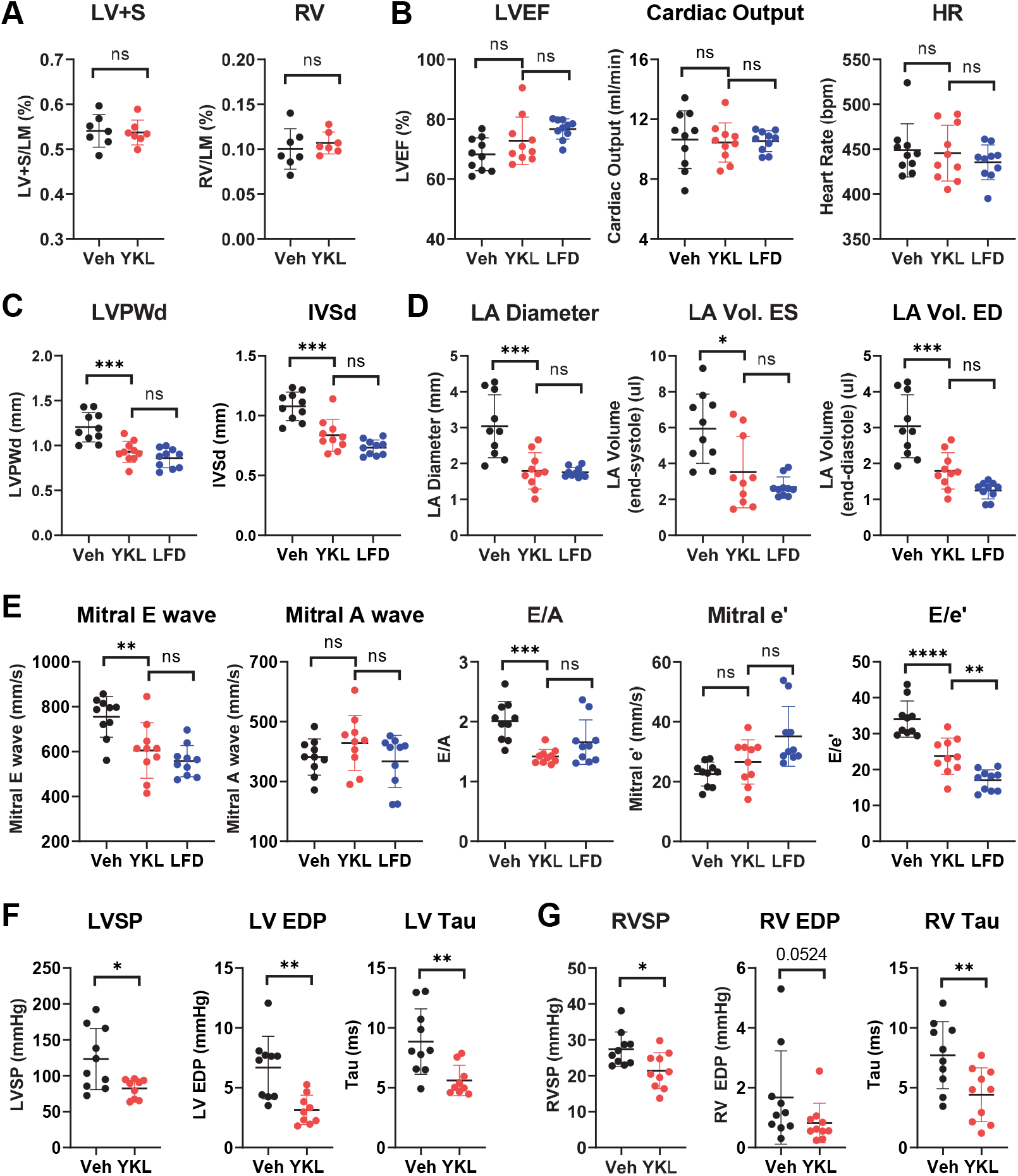
SIK inhibition improves diastolic function in obese HFpEF mice. (A) Ratio of left ventricle + septum (LV+S) and right ventricle (RV) weight to total lean mass. (B-D) Echocardiographic analyses of LV ejection fraction (LVEF), cardiac output, heart rate, end-diastolic left ventricular posterior wall thickness (LVPWd), interventricular septum thickness (IVSd), left atrial (LA) diameter, LA stroke volume end-systolic, LA stroke volume end-diastolic. (E) Doppler analysis of mitral velocity. E: early mitral wave velocity. A: late mitral wave wave velocity. e’: early diastolic mitral annular velocity. E/A: early mitral wave velocity to late mitral wave peak velocity ratio. E/e’: early mitral wave velocity to early diastolic mitral annular velocity ratio. (F-G) Pressure-volume loop measurement of LV and RV systolic pressure (SP) and end-diastolic pressure (EDP) and diastolic time constant Tau. P-values of t-test (A); ANOVA or Kruskal-Wallis test (B-E); t-test or Mann-Whitney test (F-G). *p<0.05, **p<0.01, ***p<0.001, ****p<0.0001, ns: not statistically significance.

### Dysregulation of myocardial SIK gene expression in human HFpEF

Our data suggests that SIK inhibition by YKL has a profound effect both on adiposity and cardiac function in a preclinical obese HFpEF model. However, the pathological connection of SIKs with human HFpEF is yet to be established. To this end, we analyzed a previously published myocardial transcriptomic data from healthy donor and patients with HFpEF^9^. Our results show that SIK2 and SIK3 are more abundantly expressed than SIK1 in human myocardium (**Figure 4A**). In addition, the mRNA levels of all three SIK isoforms are moderately down-regulated in myocardium of HFpEF patients (**Figure 4A**). Due to the heterogeneity of human HFpEF, three subgroups of HFpEF patients with distinct myocardial transcriptomic signatures and survival outcomes have been identified by non-negative matrix factorization and weighted gene co-expression analysis, with group 1 HFpEF patients having a greater risk of HF hospitalization and/or death^9^. Interestingly, myocardial expression of SIK1, SIK2, and SIK3 are higher in group I HFpEF patients than the other two groups (**Figure 4B**). These results indicate that myocardial SIK gene expression is dysregulated and might account for the heterogeneity and event-free survival in human HFpEF.

**Figure 4.**
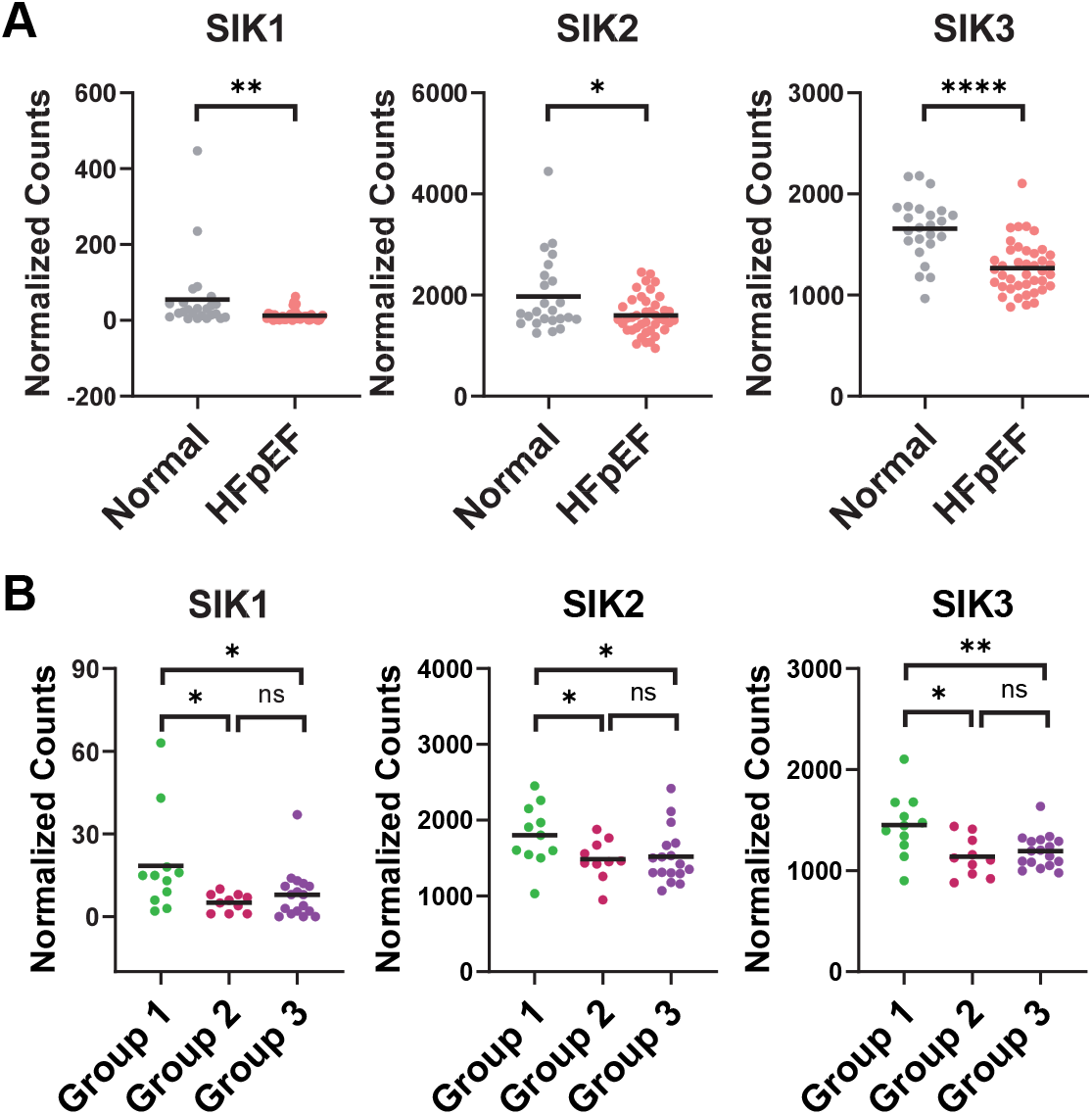
Dysregulated myocardial SIK mRNA expression in human HFpEF. (A) Myocardial *SIK1, SIK2*, and *SIK3* mRNA levels in healthy individuals and patients with HFpEF. (B) Myocardial *SIK1, SIK2*, and *SIK3* mRNA levels across three HFpEF subgroups. P-values of Mann-Whitney test. *p<0.05, **p<0.01, ***p<0.001, ****p<0.0001, ns: not statistically significance.

## Methods

### Reagents

YKL-05-099 (YKL) was purchased from MedChemExpress (HY-101147) (Monmouth Junction, NJ) or TargetMol (T17271) (Boston, MA) and dissolved in DMSO (100 mg/ml) as stock at -80C. For injection, YKL stock was reconstituted to a final concentration of 2 mg/ml in a vehicle solution that contains 5% DMSO, 40% PEG300, 5% Tween-80, 50% Saline. Nω-nitro-L-arginine methyl ester (L-NAME) hydrochloride was purchased from Sigma (N5751, St. Louis, MO) and prepared in drinking water (MillQ).

### Mouse study

All animal procedures were approved by the Institutional Animal Care and Use Committee of Vanderbilt University Medical Center and in accordance with the NIH Guide for the Care and Use of Laboratory Animals. To induce the HFpEF phenotype, eight-week-old male C57BL/6J mice (Jackson Laboratories, Bar Harbor, ME) were given L-NAME (0.5 g/L, pH 7.0, changed twice a week) in drinking water and fed with high-fat diet (60% kCal from fat, Research Diet, D12492) for 5 weeks. HFpEF mice were administered with vehicle or YKL (10 mg/kg) by daily intraperitoneal (IP) injection for 4 weeks from week 2 to 5 of L-NAME/HFD treatment. For control mice, eight-week-old male C57BL/6J mice were given drinking water and fed with low-fat chaw diet (LFD) (4.5% kCal from fat, 5LOD) for 5 weeks and injected with vehicle in the last 4 weeks. Mice were housed in group. Daily food intake and feces were manually measured and collected as per cage with free access to food and water. Fecal calorie contents were measured using 6200 Isoperibol Calorimeter (Parr) at University of Michigan Animal Phenotyping Core as previously described^22^. Glucose tolerance test was performed as previously described^23^. Body composition was measured by a minispec rodent NMR system (Brucker).

### Echocardiography and *in vivo* hemodynamics

Mouse echocardiography and cardiac catheterization were performed as described previously^24^. For echocardiography, mice were anesthetized with 2% to 3% isoflurane, applied with depilatory cream to the thorax, and then placed on a heated table in supine position. Images were acquired using the VisualSonics VEVO Imaging Systems (FUJIFILM) in B mode, M mode, and Doppler mode. Parasternal long-axis, short-axis, and modified right ventricle (RV)-centric views were obtained as described^24^. Thereafter, mice were allowed to recover for 24 hours before catheterization. For open-chest catheterization, mice were anesthetized with 2% to 3% isoflurane, orotracheally intubated with 22 g catheter, mechanically ventilated at 18 cm^3^/kg, and anesthetized with 2% to 3% vaporized isoflurane for general anesthesia. On a heated surgical table in supine, ventral-side up position, a vertical incision was made along the linea alba in the rectus abdominus sheath with scissors and cautery and extended laterally. Cautery was then used to take down the diaphragm and expose the thorax. A 1.4F Mikro-tip catheter was directly inserted into the left ventricle for measurement of pressure and volume within the left ventricle. Afterward, the catheter was removed and directly inserted into the RV. Hemostasis of the left ventricle was ensured before catheterization of the RV to ensure no effect of volume loss on findings. Hemodynamics were continuously recorded with a Millar MPVS-300 unit coupled to a Powerlab 8-SP analog-to-digital converter acquired at 1000 Hz and captured to a Macintosh G4 (Millar Instruments, Houston, TX). Volume was measured in relative volume units based on recommended calibrations from Millar Instruments. Thereafter, direct cardiac puncture was used to phlebotomize mice using a heparinized syringe, and mice were euthanized with cervical dislocation. Tissues were then harvested for downstream analyses and morphometric measurements.

### Histology

Tissues were fixed in 10% neutral buffer formalin for 24-48 hours at room temperature and then processed for embedding, paraffin section, hematoxylin and eosin (H&E) and immunohistochemistry (IHC) staining at Vanderbilt University Medical Center Translational Pathology Shared Resource (TPSR) histology core lab. IHC staining was done on inguinal white adipose tissue paraffin sections (n=3) with an anti-UCP1 antibody (1: 3000 dilution, Abcam, ab10983). To quantify lipid content, four random snapshot images were taken from the H&E-stained liver and brown adipose tissue paraffin sections (n=3) and analyzed using the positive pixel counting tool in the ImageScope software (Leica).

### Human myocardial transcriptomics

A published human myocardial bulk RNA-seq dataset from health donor and patient with HFpEF^9^ was analyzed for myocardial SIK gene expression. Three subgroups of HFpEF with distinct transcriptomic signatures were identified by non-negative matrix factorization and weighted gene co-expression analysis as described^9^.

### Statistics

Data were presented as mean ± SE M and analyzed in GraphPad Prism using Student’s t-test, ANOVA, Mann-Whitney test, and Kruskal-Wallis test. Statistical significance was defined as p < 0.05.

## Discussion

SIKs are important regulators of fuel metabolism^13-15^ and cardiovascular function^25,26^. Small molecule SIK inhibitors have been developed and have shown therapeutic potential in multiple disease areas^26,27^. Our data show that pharmacological SIK inhibition by a pan-SIKi promotes weight loss and improves diastolic function in a preclinical mouse model of obesity-related HFpEF. YKL treatment has profound effects both on cardiac function and adiposity. In the heart, YKL treatment leads to morphological and functional alterations indicative of improved LV diastolic function in obese HFpEF heart, including reductions in LV filling pressure and diastolic stiffness. In addition, YKL-treated mice show morphological changes in the left atrial and hemodynamic improvements in the right ventricle. In the YKL-treated mice, several echocardiographic parameters, including LVPWd, IVSd, LA diameter, and E/A, are reverted to an extent that is comparable to the LFD control mice, suggesting a direct disease-modifying effect in obese HFpEF hearts. As diastolic dysfunction is a major pathophysiology feature in HFpEF^5^, the treatment effects of YKL are expected to improve the cardiac outcomes in obese HFpEF mouse model.

The complexity of HFpEF pathophysiology is manifested by the presence of multiple comorbidities. About 45% of patients with HFpEF have diabetes while more than 50% are obese^28,29^. Drugs originally developed for diabetes and obesity, such as sodium-glucose transport protein 2 inhibitors and incretin-based weight loss medications, have shown clinically significant benefits in HFpEF^30-32^, suggesting targeting such comorbidities is a valid therapy for HFpEF. Our data shows that YKL treatment has a major weight loss effect. YKL treatment limits the gain of body fat, treads to reduce the fecal calorie contents, but does not affect food intake. The negative energy balance in YKL treated mice is potentially led by enhanced energy expenditure, as indicated by the induction of the key thermogenic protein UCP1 in the adipose tissue. It has been well-documented that thermogenic fat is associated with a healthier cardiometabolic profile in human^33^. Adipose tissue can contribute to cardiac function through adipocyte-derived cardiac modulating factors^34,35^. It is possible that YKL treatment can improve cardiac function potentially through indirect mechanisms by enhancing adipose tissue browning and reducing adiposity. YKL treatment also leads to a moderate improvement in glucose homeostasis. It has been documented that SIKs can control glucose homeostasis in multiple tissues through various metabolic pathways^13-15^. It is yet to be determined the tissue specific effects of YKL on glucose homeostasis and insulin sensitivity in obese HFpEF mice. Therefore, further studies are needed to characterize the impact of YKL on systemic energy metabolism and glucose homeostasis through comprehensive metabolic phenotyping analyses in future.

Previous documentation has clearly suggested the heterogeneity of human HFpEF^36^. Through non-negative matrix factorization and weighted gene co-expression analysis, three subgroups of HFpEF patients with distinct myocardial transcriptomic signatures have been identified in a previous myocardial transcriptomic study of human HFpEF^9^. Our data show that myocardial expression of SIKs is higher in a HFpEF subgroup that has a greater risk for HF hospitalization and/or death. These results underscore the heterogeneity of human HFpEF and suggest that higher myocardial SIK gene expression is possibly associated with worsened outcomes in human HFpEF. Pharmacological SIK inhibition might be more efficacious in those HFpEF patients with higher risk of hospitalization and mortality. Therefore, identifying biomarkers of SIK activity would help develop strategy to screen for the most vulnerable HFpEF population for targeted therapy.

Using a pharmacological and potentially translatable approach, our study shows the therapeutic potential of SIK inhibitor in treating obesity and HFpEF. Major progress has been made in recent years in the development of small molecule SIK inhibitors. Several SIK inhibitors have been advanced to clinical trials for the treatment of inflammatory disease and several types of cancer^26^. Such progress underscores the value of SIKs as a promising therapeutic target. With proof of efficacy, those investigational new drugs can be indicated for additional disease, such as obesity and HFpEF, in the future.

## Acknowledgement

This work was supported by American Heart Association Career Development Award 23CDA1048341 (F.S.) and Vanderbilt Cardiovascular Scientific Progress and Research Connections (C-SPaRC) Pilot Grant (S.C., V.A., F.S.). V.A. was supported by a Department of Veterans Affairs Career Development Award IK2-BX005828. S.C. was supported by National Institutes of Health grant R01 DK132236.

